# The p38MAPK-MK2-HSP27 Pathway Regulates the mRNA Stability of the Senescence-Associated Secretory Phenotype

**DOI:** 10.1101/664755

**Authors:** Hayley R. Moore, Elise Alspach, Jeffrey L. Hirsch, Joseph Monahan, Sheila A. Stewart

## Abstract

Age is a significant risk factor for the development of cancer. Both age-dependent accumulation of cell autonomous mutations within preneoplastic cells and increases in senescent stromal cells within the tumor microenvironment are thought to collaborate to drive tumorigenesis. Senescent cells express pro-tumorigenic factors termed the senescence-associated secretory phenotype (SASP), subject to a variety of regulatory mechanisms that are not fully elucidated. Previous work demonstrated that p38 mitogen-activated protein kinase (p38MAPK)-dependent regulation of AUF1 occupancy on SASP factor mRNAs post-transcriptionally stabilizes many SASP mRNAs and contributes to their increased expression. Here, we address the mechanism by which p38MAPK regulates AUF1’s occupancy and activity on SASP factor mRNAs. We found that the p38MAPK-MK2-HSP27 pathway regulates both mRNA stability and AUF1 occupancy in cells induced to senesce. Furthermore, the tumor-promoting abilities of senescent stromal cells were lost upon inhibition of MK2, suggesting that this pathway is a viable therapeutic target within the tumor microenvironment.

## Introduction

Cellular senescence arises in response to a wide array of cytotoxic stresses including replication-driven telomere attrition, DNA damage, oncogene activation and increases in ROS, and other cellular stresses. Senescence is characterized by an irreversible cell cycle arrest that is mediated by activation of p21, and subsequently p16^INK4a^ and Rb. As such, senescence functions as a potent tumor suppressor mechanism when it occurs within incipient tumor cells [1]. In addition to permanent growth arrest, senescence is characterized by an altered cell morphology that includes a flattened appearance and development of stress fibers, increased expression of senescence-associated β-galactosidase (SA-β-gal), and expression of the senescence-associated secretory phenotype or SASP (also known as the senescence messaging secretome, or SMS) [2–4]. This secretory phenotype can lead to an immune-mediated clearance of tumor cells, but when it occurs in stromal cells it functions as a potent tumor promoter [2, 5–7]. The SASP consists of a large number of pro-inflammatory cytokines and other immune modulators, matrix remodeling proteins, and pro-proliferative factors, among others, that are coordinately upregulated in senescent cells. Senescent fibroblast-derived SASP can promote epithelial cell proliferation and epithelial-to-mesenchymal transition (EMT), to allow for tumor extravasation and metastasis. In addition, the SASP can remodel the stromal compartment to create an immunosuppressive environment conducive to tumor proliferation [2, 7–9].

Immediately following a senescence-inducing stimulus, cells (referred to here as pre-senescent) begin to upregulate numerous SASP factors through the activity of transcription factors like NFκB but do not yet display classic signs of senescence, including a flattened morphology or SA-β-Gal activity [10–12]. We recently demonstrated that the maintenance of SASP factor upregulation occurs *via* a transition from a transcriptionally-driven process in pre-senescent cells to a post-transcriptional stabilization mechanism in cells displaying the morphological changes and SA-β-Gal activity characteristic of senescent cells [10]. In spite of their increased transcriptional levels, several SASP factor mRNAs in pre-senescent cells, including IL-6, IL-8, and GM-CSF, are targeted for post-transcriptional degradation by AU-rich elements (AREs) in their 3’ untranslated regions (UTRs), through a process termed ARE-mediated decay (AMD), which is driven by the ARE binding protein 1 (AUF1, also called heterogeneous nuclear ribonucleoprotein D0 or hnRNPD). However, once senescence is established as evidenced by SA-β-Gal expression, AUF1 binding is decreased and SASP mRNAs are stabilized. Elevated SASP mRNA levels are sustained predominantly by a p38MAPK-dependent mRNA stabilization mechanism. Indeed, p38MAPK inhibition with the small molecule inhibitor SB203580 prevented the removal of AUF1 from these 3’UTRs and consequently the mRNA stabilization phenotype [10].

During AMD, AUF1 binds to target AREs and recruits other *trans*-acting factors (termed the AUF1- and signal transduction-regulated complex or ASTRC), which results in mRNA degradation. The precise mechanisms and order of events that lead to mRNA degradation remain to be determined, but it is thought to occur through acceleration of deadenylation and subsequent nucleolytic degradation of the target mRNA [13, 14]. While AUF1 is known to be important in this process, its precise role in mRNA decay is incompletely understood. AUF1 is expressed in four splice isoforms generated from a common pre-mRNA: p37, p40, p42, and p45. Although all isoforms can promote mRNA degradation, there are circumstances wherein certain isoforms of AUF1 bind and act to protect mRNAs from decay, underscoring the complexity of the system and demonstrating that AUF1-mediated decay is not regulated simply at the level of AUF1 binding [15–18]. Post-translational modification of AUF1 also appears to differentially affect mRNA stability. p40^AUF1^ can be phosphorylated on Ser^83^ by protein kinase A (PKA) and Ser^87^ by glycogen synthase kinase 3β (GSK3β), and this phosphorylation has contradictory consequences on mRNA stability [19, 20]. In THP-1 monocytic leukemia cells, phosphorylated p40^AUF1^ is actively involved in mRNA degradation. Treatment with 12-*O*-tetradecanoylphorbol-13-acetate (TPA) to simulate monocyte adherence results in rapid stabilization of the target mRNAs and concurrent loss of p40^AUF1^ phosphorylation, suggesting that p40^AUF1^ promotes mRNA degradation when phosphorylated, and mRNA stabilization in its non-phosphorylated state [20]. In contrast, phosphorylation of AUF1 leads to its ubiquitin-mediated degradation (and subsequent AUF1 target mRNA stabilization) in response to p38MAPK signaling through HSP27 in HeLa cells [21]. The varied consequences of AUF1 phosphorylation and differential outcomes of AUF1 binding to target AREs demonstrate that complex regulatory mechanisms including the phosphorylation state of AUF1 and/or other ASTRC members are involved in AMD.

We previously demonstrated that p38MAPK inhibition led to destabilization of SASP mRNAs through alteration of AUF1 binding in senescent fibroblasts. However, p38MAPK does not directly target AUF1 and thus the mechanism by which p38MAPK influences AUF1 activity remained to be elucidated. Interestingly, recent work by Herranz *et al.* demonstrated that MAPKAPK2 (MK2) in senescent IMR90 lung fibroblasts participated in SASP factor regulation. MK2 is a p38MAPK target which when activated is known to regulate mRNA stability and phosphorylate the small heat shock protein HSP27, an ASTRC member that has been shown to regulate AUF1 levels and mRNA stability in lymphocytes [22–25].

Given our previous findings, we investigated whether p38MAPK regulates mRNA stabilization in senescent cells through MK2 and HSP27. Our results indicate that p38MAPK and MK2 govern mRNA stabilization through phosphorylation of HSP27, but that HSP27 does not act solely through regulation of AUF1 protein levels or binding to SASP mRNAs. Furthermore, we find that depletion or inhibition of MK2 not only prevents the upregulation and stabilization of SASP factor mRNAs, it also prevents senescent fibroblasts from promoting preneoplastic cell growth in a co-culture model, suggesting that targeting MK2 in the tumor microenvironment is a promising therapeutic avenue in cancer treatment.

## Materials and Methods

### Cell lines and treatments

BJ human foreskin fibroblasts were cultured in DMEM supplemented with 15% M-199, 15% heat-inactivated FBS, and 1% penicillin/streptomycin (all from Sigma Aldrich, St. Louis, MO). Fibroblasts were treated with bleomycin sulfate (0.1 units/mL, Sigma Aldrich, St. Louis, MO) for 24 hours, followed by incubation in normal culture medium (unless otherwise stated) for the time points indicated. Fibroblasts were treated with actinomycin D (10 μg/mL, Sigma Aldrich, St. Louis, MO) for 24 hours, and SB203580 (10 μM, Millipore, Billerica, MA), CDD-111 (also referred to as SD0006, 1 μM), or CDD-450 (1μM, both from Confluence Life Sciences, St. Louis, MO) every 24h unless indicated otherwise. HaCaT preneoplastic keratinocyte cells obtained from Dr. Norbert E. Fusenig (German Cancer Research Center, Heidelberg, Germany) stably expressing click beetle red (CBR) luciferase (HaCaT-CBR) and HEK293T embryonic kidney cells [6] were grown in DMEM supplemented with 10% heat-inactivated FBS and 1% penicillin/streptomycin (Sigma Aldrich, St. Louis, MO). All cells were cultured at 37°C in 5% CO_2_ and 5% O_2_. No cell lines used were authenticated.

### Virus production and plasmids

Virus production was carried out as described previously [26]. Briefly, HEK239T cells were transfected with the target plasmid, pCMV-VSV-G, and either pHR’-CMV-8.2ΔR for pLKO.1 and pRESQ constructs, or pUMVC3 for pBABE constructs, using Trans-IT LT1 (Mirus, Madison, WI) and virus was collected 48h later. Infections were carried out in the presence of 1μg/mL protamine sulfate. 48h post-infection, cells were selected with 1μg/mL puromycin or 50 μg/mL hygromycin.

Short hairpin RNA sequences targeting HSP27 (5’-CCCGGACGAGCTGACGGTCAA-3’), MK2 (5’-AGAAAGAGAAGCATCCGAAAT-3’) and control SCR (5’-TCCTAAGGTTAAGTCGCCCTC-3’) were obtained from the Children’s Discovery Institute’s viral vector-based RNAi core at Washington University in St. Louis, and were supplied in the pLKO.1-puro backbone. When not combined with knockdown, FLAG-tagged HSP27-TriD was expressed from the pBABE-hyrgo backbone. Knockdown rescue experiments utilized the pRESQ-puro backbone with the indicated hairpins. Hairpin-resistant Flag-HSP27 WT and TriD were manufactured by IDT (Coralville, IA) and cloned into pRESQ from pUC57. The sequence was based on constructs provided by Dr. Gary Brewer with the following silent mutations to the shHSP27 hairpin recognition sequence: 5’-CC<u>G</u> GA<u>C</u> GA<u>G</u> CT<u>G</u> AC<u>G</u> GT<u>C</u>-3’ to 5’-CC<u>C</u> GA<u>T</u> GA<u>A</u> CT<u>C</u> AC<u>C</u> GT<u>G</u>-3’. Hairpin-resistant HSP27-TriA was generated utilizing QuikChange Lightning site directed mutagenesis (Agilent Technologies, Santa Clara, CA) of pUC57-HSP27-WT and published primers [23] to generate the Ser-to-Ala mutations at serines 15, 78 and 82, and similarly cloned into pRESQ.

### Western blot analysis

Cell pellets were lysed in MCLB (50 mM Tris pH 8.0, 5 mM EDTA, 0.5% NP40 and 100 mM sodium chloride, with aprotenin, leupeptin, pepstatin, phenylmethylsulfonyl fluoride, microcystin LR, sodium orthovanidate and sodium fluoride) for 20 minutes at 4°C. Protein concentration was quantified using the Bradford Protein Assay (Bio-Rad, Berkeley, CA). The primary antibodies used were: p-HSP27 Ser^82^ (1:1000, catalog number 2401S), p-HSP27 Ser^15^ (1:1000, catalog number 2404S), monoclonal HSP27 (1:1500, catalog number 2402S), MK2 (1:1000, catalog number 3042) and p38 (1:1000, catalog number 9218S) all from Cell Signaling Technology (Danvers, MA); p-HSP27 Ser^78^ (1:2000, clone Y175, catalog number ab32501, Abcam, Cambridge, MA); AUF1 (1:4000, catalog number 07-260, Millipore); p-p38 (1:1000, catalog number p190-1802, PhosphoSolutions, Aurora, CO); α-tubulin (1:5000, catalog number ab6160, Abcam, Cambridge, MA); γ-actin (1:5000, catalog number NB600-533, Novus Biologicals, Littleton, CO); β-actin (1:2000, clone AC-15, catalog number A1978) and GAPDH (1:2500, clone GAPDH-71.1, catalog number G8795) both from Sigma Aldrich, St. Louis, MO. Additional primary antibody information can be found in S1 Table. All secondary antibodies from the appropriate species were horseradish peroxidase-conjugated (Jackson Laboratories, Bar Harbor, ME) and diluted 1:10000. Western blots were imaged using a Bio-Rad Chemidoc XRS+ imager and ImageLab software (Version 3.0 build 11, Berkeley, CA).

### Quantitative PCR

RNA was isolated using TRI Reagent (Life Technologies, Carlsbad, CA) or RNeasy kit (Qiagen, Hilden, Germany) at the time points indicated. cDNA synthesis and quantitative PCR was performed using previously published protocols and manufacturers’ instructions [10] (SYBR Green, Life Technologies, Carlsbad, CA). Primers for GAPDH (F: 5’-GCATGGCCTTCGGTGTCC-3’, R: 5’-AATGCCAGCCCCAGCGTCAAA-3’), IL-6 (F: 5’-ACATCCTCGACGGCATCTCA-3’, R: 5’-TCACCAGGCAAGTCTCCTCA-3’), IL-8 (F: 5’-GCTCTGTGTGAAGGTGCAGT-3’, R: 5’-TGCACCCAGTTTTCCTTGGG-3’), GMCSF cDNA was amplified using a Taqman probe/primer set and normalized to GAPDH (catalog numbers Hs00929873_m1 and Hs02758991_g1, respectively, and Taqman Fast Advanced master mix, Life Technologies, Carlsbad, CA).

### p38MAPK enzyme activity assay

CDD-450 and CDD-110 were synthesized by Confluence Discovery Technologies and provided as solid powders by the Medicinal Chemistry Department. CDD-110 is a prototypic global p38 inhibitor, which is the racemate of PH-797804 [27]. Compounds were dissolved in 100% DMSO (J. T. Baker, Aldrich Chemical, St. Louis, MO) to a concentration of 10 mM.

### *In vitro* activation of p38MAPKα

Nonactive p38MAPKα was phosphorylated and activated *in vitro* prior to use in the kinase assays described below. Nonactive p38MAPKα (2 mM) was incubated with 200 mM MgATP and 40 nM constitutively active MKK6 in kinase buffer (20 mM HEPES pH 7.6, 10 mM MgCl_2_, 0.01% TritonX-100, 0.01% BSA, 1 mM DTT) for 2 hours at room temperature. After the incubation period, aliquots of activated p38MAPKα were stored at −80°C for use in kinase assays.

### p38MAPKα/MK2 Kinase Cascade Assay

An assay was developed that quantifies the activity of p38MAPKα toward MK2 by its ability to phosphorylate and activate the downstream kinase. The kinase activity of phosphorylated MK2 is followed by measuring its phosphorylation of a fluorescently-labeled, peptide substrate based on the phosphorylation sequence of the physiological substrate of MK2, Hsp27 (FITC-KKKALSRQLSVAA, American Peptide catalog number 310945, Sunnyvale, CA). The phosphorylation of the Hsp27 peptide was quantified using IMAP Progressive Screening Express Kit (Molecular Devices, catalog number R8127, Sunnyvale, CA) and the Analyst HT plate reader. Kinase reactions are carried out in a 384-well assay plate in kinase buffer with a final DMSO concentration of 2%. CDD-450 and CDD-110 were tested in duplicate at 11 concentrations using a 3-fold serial dilution scheme. Hsp27 peptide substrate and MgATP were held constant at 1 μM and 10 μM, respectively. Activated p38MAPKα was added to a final concentration of 60 pM for reactions with 1nM non-phosphorylated GST-MK2 (Millipore Corporation, catalog number 14-349, Billerica, MA). Kinase reactions were incubated at room temperature and quenched after 120 minutes by the addition of 1X Progressive Binding Buffer A containing 1X IMAP beads. After addition of IMAP stop solution, reaction mixture was incubated at room temperature for 1 hour to allow binding of the phosphorylated peptide product to the IMAP beads to come to equilibrium. The assay plate was then read on Analyst HT plate reader using standard fluorescence polarization settings. Inhibitor potency was calculated by fitting dose-response data to the 4-parameter logistical IC_50_ equation.

### p38MAPKα/PRAK Kinase Cascade Assay

An assay was developed that quantifies the activity of p38MAPKα toward its substrate PRAK by its ability to phosphorylate and activate the downstream kinase. The kinase activity of phosphorylated PRAK was followed by measuring the phosphorylation of the fluorescently-labeled peptide substrate HSP27, as described above. Kinase reactions were carried out in a 384-well assay plate in kinase buffer with a final DMSO concentration of 2% and contained 1 μM Hsp27 peptide and 10 μM MgATP. CDD-450 and CDD-110 were tested in duplicate at 11 concentrations using a 3-fold serial dilution scheme. Activated p38MAPKα was added to a final concentration of 62.5 pM for reactions with 10nM non-phosphorylated GST-PRAK. Kinase reactions were incubated at room temperature and quenched after 120 minutes by the addition of 1X Progressive Binding Buffer A containing 1X IMAP beads. After quenched reactions equilibrate for 1 hour, the assay plate is read and data is calculated as described above.

### Ribonucleoprotein (RNP) immunoprecipitation (RIP)

Cell pellets from two 150mm dishes of BJ fibroblasts were lysed in the same buffer used for western blot analysis (MCLB). Protein concentration was analyzed using the Bradford Protein Assay (Bio-Rad, Berkeley, CA). 1 mg of protein was used for each immunoprecipitation. 15 μg of polyclonal AUF1 (catalog number 07-260, Millipore, Billerica, MA) was used. An equivalent amount of normal rabbit IgG antibody (catalog number 2729S, Cell Signaling, Danvers, MA) was used to control for specific immunoprecipitation. Cell lysates were pre-cleared with 20 μL protein A Dynabeads (Life Technologies, Carlsbad, CA) for 30 minutes at 4°C prior to incubation with the indicated antibody overnight at 4°C in total volume of 1mL MCLB supplemented with 1 μL RNAseOUT (Invitrogen, Carlsbad, CA). 100 μL Protein A Dynabeads were used for each immunoprecipitation. Beads were washed 2 times in 0.1 M monosodium phosphate, 3 times in Buffer A (1x PBS, 0.1% SDS, 0.3% sodium deoxycholate, 0.3% NP40), and 2 times in MCLB, then incubated with samples for 4 hours at 4°C with rotation. Immunoprecipitated beads were washed 2 times with each of the following buffers: Buffer A, Buffer B (5x PBS, 0.1% SDS, 0.5% sodium deoxycholate, 0.5% NP40) and Buffer C (50 mM Tris pH 7.4, 10 mM magnesium chloride, 0.5% NP40). RNA was isolated from the beads by adding 1 mL of TRI Reagent (Life Technologies, Carlsbad, CA). Following cDNA synthesis, mRNA levels of SASP factors were analyzed by qPCR using the primers and procedures described above.

### Senescence-associated β-galactosidase

SA-β-gal staining was carried out on subconfluent cells as described previously [6, 28]. Briefly, sub-confluent cells were fixed in 0.2% glutaraldehyde for 5’ at room temperature, washed in PBS, and incubated at 37°C in X-gal staining solution (40mM NaPi pH 6.0, 5 mM K_4_Fe(CN)_6_, 5mM K_3_Fe(CN)_6_, 2mM MgCl_2_, 1 mg/mL X-Gal (5-Bromo-4-chloro-3-indolyl-β-D-galactopyranoside, Sigma Aldrich Aldrich, St. Louis, MO), and 150mM NaCl, and 1mg/mL) until color was sufficiently developed (∼4h). Staining was stopped by washing 2X with PBS, and images were acquired using a Nikon Eclipse T*i*-S (Nikon Instruments, Melville, NY) with Retiga EXi-B camera and Q-capture 64 software (QImaging, British Columbia, Canada).

### Co-culture

Co-culture experiments were performed as previously described [10]. Briefly, 1.3×10^4^ BJ fibroblasts were plated in black-walled 96-well plates. 24h later, cells were treated with 0.1 units/mL bleomycin for 24h. Cells were then incubated in starve medium (DMEM F-12 + 1% penicillin/streptomycin) for 24h before addition of CDD-111 and CDD-450. Inhibitors were refreshed daily for 2 days before the addition of HaCaT-CBR cells. HaCaT-CBR cells were cultured in starve medium for 24 hours prior to plating on fibroblasts. 1.0×10^3^ HaCaT-CBR cells were plated on fibroblasts and incubated with inhibitors for 72h. After 72h, D-luciferin (Biosynth, Naperville, IL) was added to a final concentration of 150 μg/mL. After ten minutes, plates were imaged using an IVIS 100 camera (PerkinElmer, Downers Grove, IL) using the following settings: exposure=5 min, field of view=15, binning=16, f/stop=1, open filter.

## Results

### HSP27 destabilizes SASP factor mRNA in pre-senescent cells

SASP factor mRNAs are regulated at the transcriptional and post-transcriptional levels [10]. Indeed, our previous work demonstrated that the stress kinase p38MAPK stabilizes SASP factor mRNAs by altering the RNA binding of AUF1, which can target these mRNAs for degradation. Importantly, the increased mRNA stability is induced only after the establishment of the senescent phenotype as indicated by expression of senescence-associated β-galactosidase (SA-β-Gal) and morphological changes consistent with the induction of senescence. In verification of our previous results, when transcription is inhibited with actinomycin D (ActD) immediately following a senescence-inducing dose of bleomycin but before senescence phenotypes such as SA-β-Gal expression and increases in p38MAPK phosphorylation are evident (pre-senescent), the abundance of the SASP factor mRNAs IL6, IL8, and GM-CSF were significantly reduced compared to similar treatment of senescent cells (Fig 1A-C). The transcriptional stabilization correlates with protein expression, as transcriptional inhibition results in decreased IL6 protein levels as measured by ELISA in pre-senescent, but not senescent cells [10]. This indicates that transcriptionally-driven mechanisms predominantly contribute to increased SASP expression in pre-senescent cells, but that the increased levels of SASP factors observed in senescent cells is due to the activation of a post-transcriptional mRNA stabilization mechanism.

**Fig 1.**
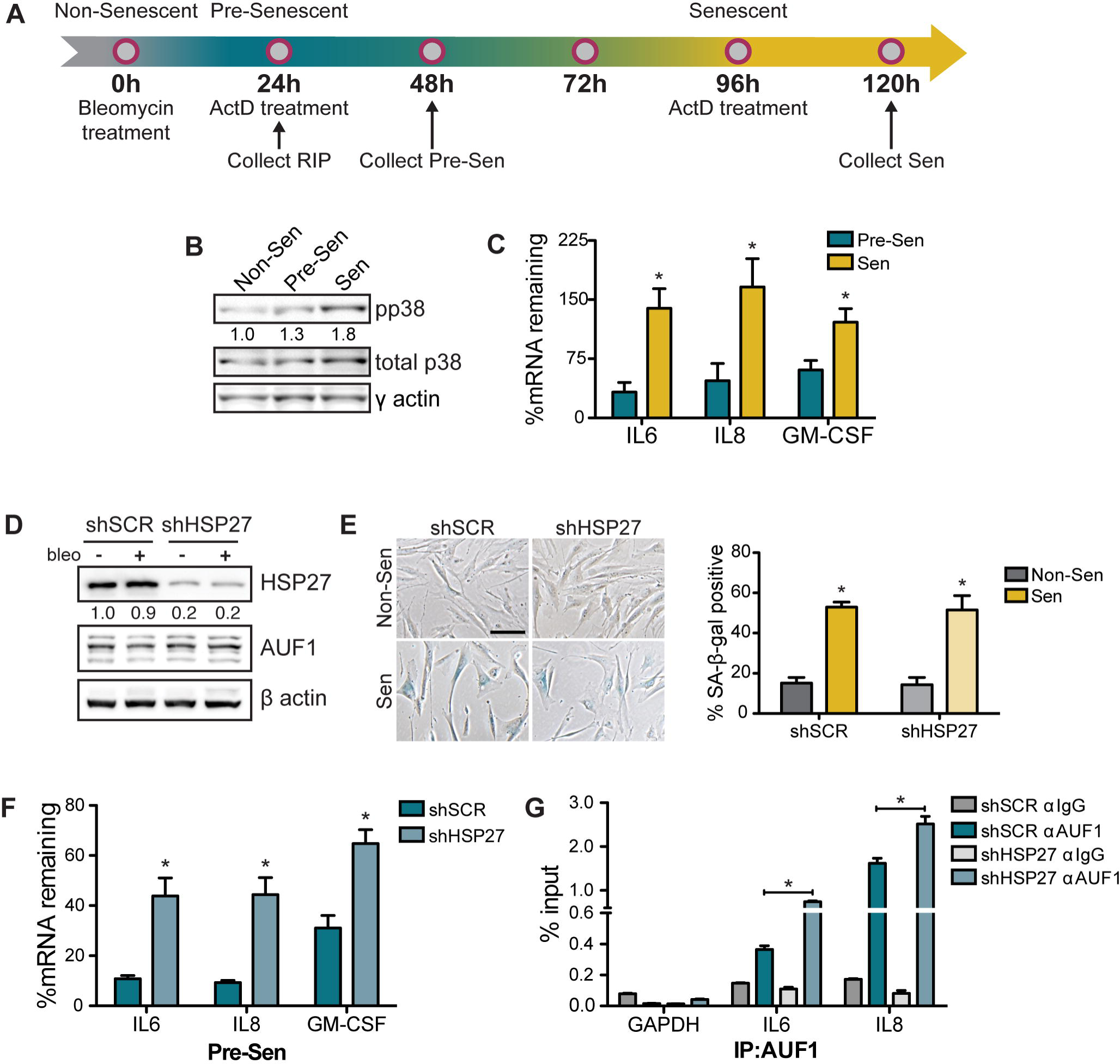
HSP27 promotes SASP mRNA degradation in pre-senescent cells. (A) Timeline of bleomycin and actinomycin D (ActD) treatment to induce stress-induced premature senescence and inhibit transcription in BJ fibroblasts. Fibroblasts were treated with bleomycin (0.1 units/mL) for 24h and subsequently treated with 7.96μM ActD for 24h starting either 24h or 96h after bleomycin treatment. Cells collected at 48h were considered pre-senescent (Pre-Sen) while those collected at 120h were considered senescent (Sen). (B) Immunoblot of phosphorylated and total p38MAPK in cells collected at the Non-Sen, Pre-Sen, and Sen points as indicated in A. Quantification of phospho-p38MAPK is relative to total p38MAPK and appears under the phosphorylated p38MAPK (pp38) blot. (C) Cells were collected after ActD treatment and SASP factor levels were quantified by qRT-PCR. Percent mRNA (%mRNA) remaining was calculated as the ratio of ActD-treated to vehicle-treated levels of target mRNA, normalized to GAPDH, n=6. (D) Immunoblot of indicated proteins in cells expressing an shSCR or shHSP27 hairpin, treated as indicated. The ratio of knockdown is reported under the HSP27 blot as amount of protein remaining in shHSP27 compared to shSCR cells after normalizing to β-actin. (E) Representative images and quantification of senescence-associated β-galactosidase (SA-β-Gal) activity in non-senescent and senescent cells expressing either shSCR or shHSP27. Images acquired with a 10x objective, scale bar = 100μm, n=3. (F) Control or HSP27-depleted cells were treated as in A, ActD treated at the pre-senescent timepoint, and collected 24h later. n=9 for IL-6 and IL-8, and n=3 for GM-CSF. (G) BJ fibroblasts expressing either shSCR or shHSP27 were induced to senesce and collected 24h post-bleomycin addition (pre-senescent), and AUF1-bound RNA was isolated by RNA immunoprecipitation. Lysates were incubated with 15 μg of either control rabbit IgG (gray bars) or rabbit anti-AUF1 antibody (blue bars). The RNA bound to precipitated protein was isolated and levels were analyzed by qRT-PCR. Representative experiment shown, n=4. * = p<0.05, error bars are + S.E.M.

Our previous work demonstrated that p38MAPK inhibition decreased SASP factor mRNA stability in senescent cells in part by allowing AUF1 to remain active and bound to the UTRs of these mRNAs, resulting in their degradation [10]. Because AUF1 is not a direct p38MAPK target, we wanted to determine how p38MAPK impacted the ability of AUF1 to regulate SASP factor mRNA degradation following senescence induction. We turned our attention to MK2 because it phosphorylates HSP27 following p38MAPK activation, and can regulate mRNA stability as a member of the AUF1-containing ASTRC complex in other experimental settings [23, 24, 29]. To determine whether HSP27 could regulate mRNA stability in pre-senescent cells, we transduced BJ fibroblasts with a control short hairpin (shSCR) or a short hairpin targeting HSP27 (shHSP27), resulting in an 80% reduction of HSP27 protein (Fig 1D). Next, we treated control and HSP27-depleted cells with bleomycin to induce senescence and found that HSP27 depletion did not significantly affect the ability of cells to enter senescence as quantified by SA-β-Gal activity (Fig 1E). 24h after bleomycin addition, cells were treated with ActD to assess the stability of SASP factor transcripts. Indeed, there was a significant increase in IL6, IL8, and GM-CSF mRNA remaining in pre-senescent cells depleted of HSP27 relative to shSCR-expressing cells after transcriptional inhibition (Fig 1F), demonstrating that HSP27 acts to destabilize target SASP mRNAs in pre-senescent cells. Previous studies have demonstrated that HSP27 can regulate AUF1 levels, suggesting that HSP27 regulates mRNA stability by modulating AUF1 abundance [21, 23]. In our system, however, we failed to find evidence that HSP27 depletion consistently modulated AUF1 abundance, suggesting the mRNA stabilization we observed did not result from changes in overall AUF1 levels (Fig 1D, data not shown).

shRNA-mediated depletion of AUF1 stabilizes SASP factor mRNAs in pre-senescent cells suggesting AUF1 functions to destabilize these mRNAs [10]. Therefore, we next wanted to determine whether HSP27 affected the ability of AUF1 to bind to SASP mRNAs and alter their stability. To test this possibility, we utilized RNAi to deplete HSP27 from BJ fibroblasts, and induced senescence via bleomycin treatment. We then performed a ribonucleoprotein (RNP) immunoprecipitation (RIP) to capture AUF1-bound mRNAs. GAPDH mRNA, which does not contain an ARE, was not significantly immunoprecipitated by an anti-AUF1 antibody relative to IgG control, but the anti-AUF1 antibody did immunoprecipitate the ARE-containing IL6 and IL8 mRNAs. Surprisingly, depletion of HSP27 resulted in increased AUF1 binding to IL6 and IL8 mRNAs, from 0.36% to 0.74% of input for IL6, and 1.62% to 2.51% of input for IL8 in shSCR versus shHSP27-expressing cells (Fig 1G). This finding demonstrates that although AUF1 is necessary for mRNA degradation in pre-senescent cells [10], its binding alone is insufficient to induce mRNA degradation in pre-senescent fibroblasts depleted of HSP27. Together these data suggest that an additional HSP27-dependent activity or HSP27-dependent alteration in the binding of another ARE binding protein is required to drive ARE-mediated decay in pre-senescent cells.

### p38MAPK and MK2 regulate mRNA stability in senescent cells

We previously demonstrated that p38MAPK regulates the transition from unstable SASP factor mRNAs in pre-senescent cells to stable mRNAs with longer half-lives in senescent cells [10]. Given HSP27’s role in SASP factor stabilization and the fact that MK2 can impact SASP expression as well as induce stabilization of ARE-containing mRNAs in response to p38MAPK activation [22, 23, 25, 30], we next interrogated the role the p38MAPK-MK2 pathway plays in the senescence-associated mRNA stabilization phenotype. To address this question, we utilized several inhibitor compounds. SB203580 (p38i) is a small molecule inhibitor of p38MAPKα and p38MAPKβ, which we used at a concentration that minimizes off-target effects (10μM) [31, 32]. CDD-450 (MK2i) is a novel p38MAPK/MK2 pathway inhibitor that selectively targets the p38MAPK/MK2 complex, inhibiting activation of MK2 by p38MAPK while allowing p38MAPK to phosphorylate other downstream targets (Fig 2). Both MK2 and a closely related downstream kinase, PRAK, require phosphorylation by p38MAPK for activation. Treatment with the p38MAPK inhibitor p38i′ (CDD-110) inhibited both MK2 and PRAK activity, as measured by decreased phosphorylation a fluorescently-labeled HSP27 peptide (Fig 2A). In contrast, treatment with MK2i inhibited MK2 activity, but not PRAK activity (Fig 2B). Therefore, MK2i provides us with a tool to selectively investigate the effects of p38MAPK-dependent MK2 activation on SASP mRNA stability. As expected, treatment with p38i or MK2i resulted in an approximately 70% reduction in p38MAPK and MK2 activity, respectively, as measured by average inhibition of HSP27 phosphorylation (Fig 3A-B).

**Fig 2.**
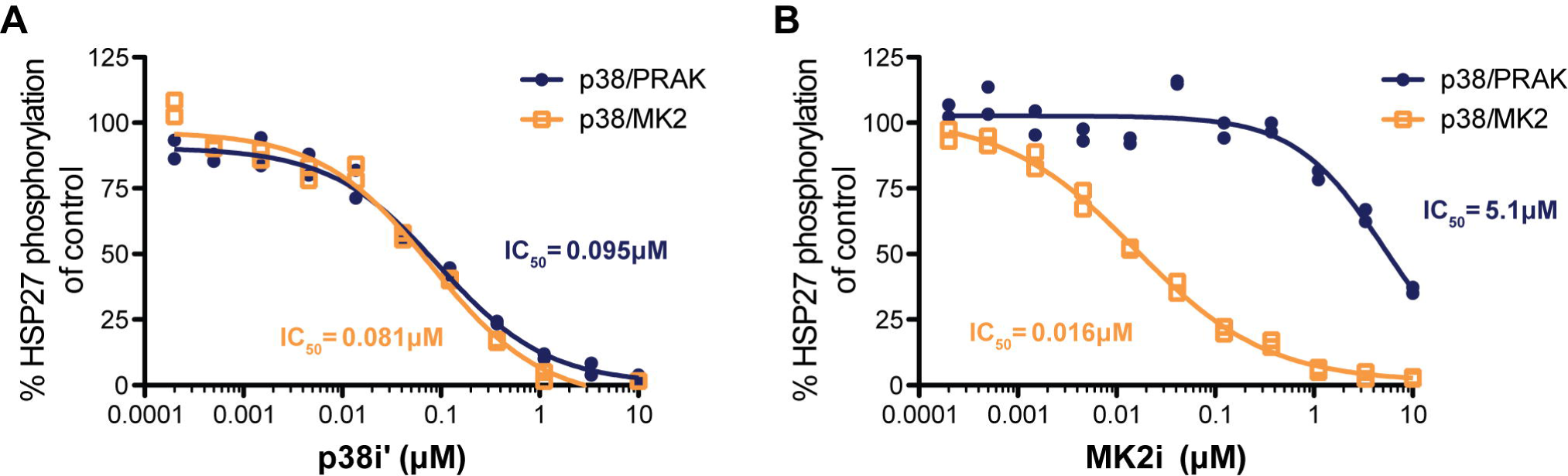
CDD-450 selectively inhibits p38MAPK-MK2 activity. The selectivity of MK2i/MK2 complex as compared to the p38i′ inhibitor CDD-110 was assayed by measuring the ability of the compounds to inhibit the kinase activities of MK2 and PRAK. Both MK2 and PRAK require p38MAPKα phosphorylation for activation. MK2 and PRAK activity was measured *via* their ability to phosphorylate a fluorescently-labeled HSP27 peptide. 1nM non-phosphorylated GST-MK2 or 10nM non-phosphorylated PRAK was incubated were 62.5pM or 60pM of activated p38α, respectively, and a 3-fold dilution series of (A) p38i or (B) MK2i. Phosphorylation of HSP27 was quantified using IMAP assay technology, and the inhibitor potency was calculated. Representative experiment, n=8.

**Fig 3.**
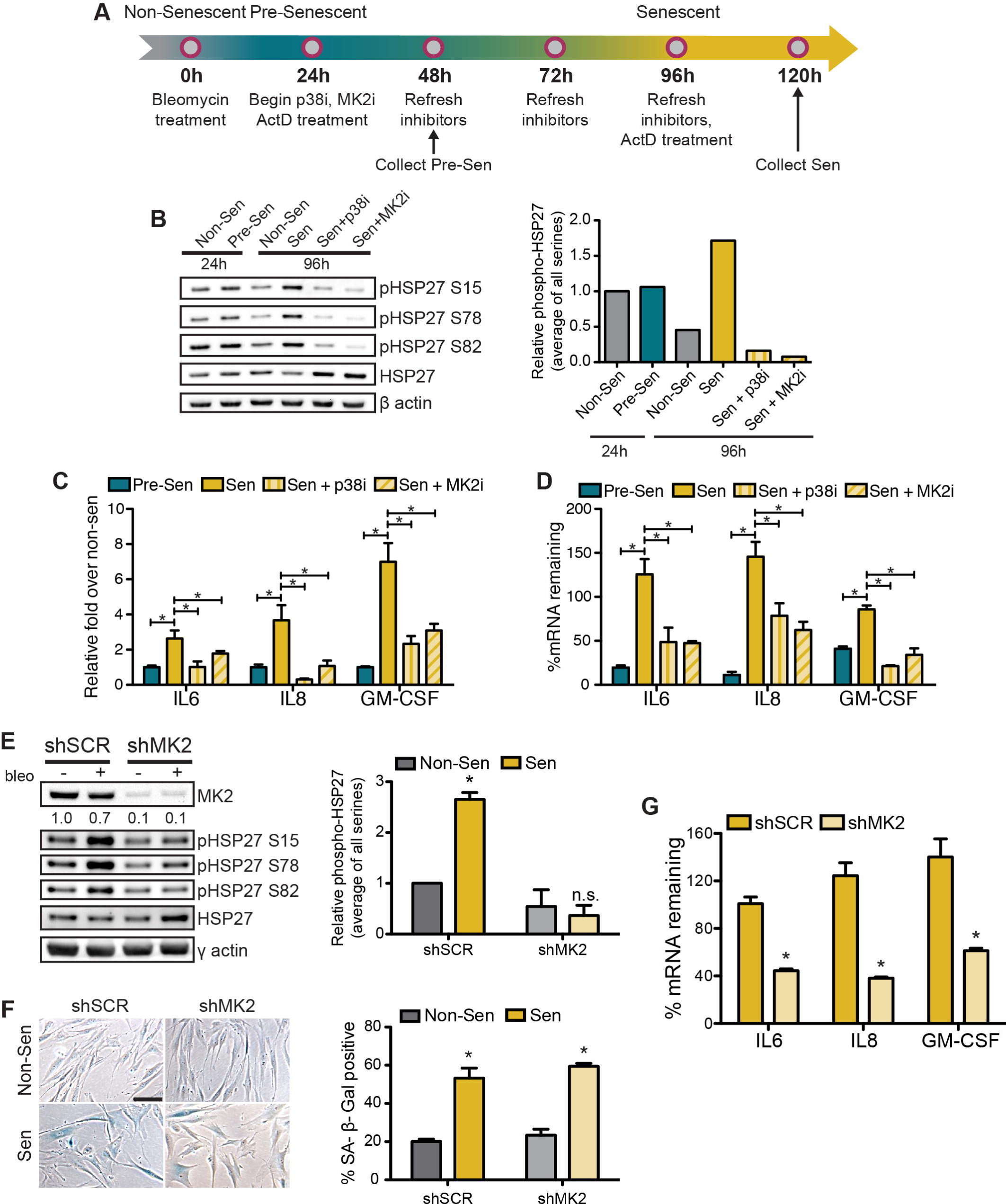
p38MAPK and MK2 regulate SASP mRNA stability in senescent cells. (A) Schematic of senescence induction by bleomycin treatment and inhibition of p38MAPK and MK2 pathways by treatment with 10 μM CDD-450, respectively. (B) Immunoblot and representative quantification (of immunoblot shown) of phospho-HSP27 at Ser15, Ser78 and Ser82, the three MK2 targets, upon p38MAPK or MK2 pathway inhibition 48h (Pre-Sen) and 120h (Sen) post-bleomycin treatment. Phospho-HSP27 was normalized to total HSP27 and the ratio of phosphorylation at each site is quantified and averaged. (C) Upregulation of SASP factors in senescent cells treated with p38i or MK2i as in A without transcriptional inhibition. mRNA levels were determined by qRT-PCR and are represented as fold increase over non-senescent cells, normalized to GAPDH. (D) Cells were induced to senesce and treated with p38i and MK2i as shown in A, and ActD treated to inhibit transcription for 24h. %mRNA remaining is calculated as the amount of mRNA in ActD-treated cells relative to vehicle-treated cells. C and D are representative experiments, n=5, *=p<0.05, error bars are ± S.E.M. (E) Immunblot and quantification of Non-Sen and Sen cells expressing shSCR or shMK2, demonstrating ∼90% knockdown of MK2 at protein level, normalized to γ actin, then to Non-Sen shSCR. Cells were induced to senesce and collected as in A. HSP27 phosphorylation is quantified and averaged. Quantification n=2. (F) Representative images and quantification of senescence-associated β-galactosidase (SA-β-Gal) activity in non-senescent and senescent cells expressing either shSCR or MK2. Images acquired with a 10x objective, scale bar = 100μm, n=2. (G) Control or MK2-depleted cells were treated as in A, ActD treated at the senescent timepoint, and collected 24h later. n=2. * = p<0.05, error bars are + S.E.M.

To confirm the role of p38MAPK activity in the transition from transcriptional regulation to post-transcriptional stabilization of SASP mRNAs as cells become senescent, we assayed mRNA stability in cells treated with p38i. As previously reported [10], we found that treatment with p38i significantly reduced the expression of the SASP factors IL6, IL8, and GM-CSF by 62%, 92%, and 67% relative to vehicle-treated senescent cells (Fig 3C). p38i also significantly inhibited the stabilization of these mRNAs in senescent cells, by 61%, 46%, and 75% for IL6, IL8, and GM-CSF, respectively (Fig 3D). Similarly, MK2i treatment of senescent cells inhibited expression of IL6, IL8, and GM-CSF by 32%, 71%, and 56% relative to vehicle-treated senescent cells (Fig 3C). Furthermore, MK2 pathway inhibition recapitulated our findings with p38i by preventing mRNA stabilization in senescent cells, with a 62%, 57%, and 60% reduction in mRNA remaining after ActD treatment for IL6, IL8, and GM-CSF, respectively (Fig 3D). Thus, MK2 activity is required for the stabilization of SASP factor transcripts in senescent fibroblasts.

While CDD-450 is highly specific for the p38MAPK-MK2 interface, we wanted to confirm that MK2 plays a critical role in SASP factor mRNA stabilization. To do this, we utilized an RNAi approach and designed an shRNA construct that specifically targeted MK2, depleting greater than 90% of MK2 at the protein level (Fig 3E). MK2 depletion did not significantly affect the ability of the fibroblasts to enter senescence as indicated by SA-β-Gal expression (Fig 3F).

If MK2 were the kinase responsible for the senescence-dependent phosphorylation of HSP27, MK2 knockdown should reduce the levels of senescence-induced HSP27 phosphorylation. Indeed, we observed a reduction in HSP27 phosphorylation at all three MK2 target serines in senescent cells expressing shMK2 compared to cells expressing shSCR (Fig 3E). To determine whether MK2 was required for stabilization of SASP mRNAs in senescent cells, we inhibited transcription in senescent cells expressing either shSCR or shMK2 and compared the levels of SASP factor mRNAs (Fig 3A, without p38 or MK2 inhibitor treatment). Knockdown of MK2 resulted in a significant reduction in the amount of SASP factor mRNA remaining after transcriptional inhibition, by 70% for IL8 and approximately 55% for IL6 and GM-CSF (Fig 3G). Together, these data demonstrate that MK2 is required for both the observed phosphorylation of HSP27 in senescent cells, and for stabilization of SASP factor mRNAs during senescence. Further, given that inhibition of p38MAPK-dependent activation of MK2 by MK2i results in similar reductions in HSP27 phosphorylation and mRNA stabilization, these data together suggest that MK2 is the downstream kinase responsible for p38MAPK-dependent SASP mRNA stabilization in senescent cells.

### The p38MAPK-MK2-AUF1 pathway regulates mRNA stability in senescent cells through HSP27 phosphorylation

Because HSP27 is required for mRNA degradation in pre-senescent cells, and p38MAPK and MK2 are required for SASP factor mRNA stability in senescent cells, we next asked if p38MAPK-MK2-dependent HSP27 phosphorylation impacted mRNA stability in pre-senescent and senescent cells, as is observed in other contexts [21, 23, 24]. Analysis of HSP27 phosphorylation revealed a p38MAPK- and MK2-dependent increase in phosphorylation at the MK2 target sites Ser^15^, Ser^78^, and Ser^82^, in senescent versus pre-senescent cells (Fig 3B). To investigate the role of p38MAPK-MK2-dependent HSP27 phosphorylation, we simultaneously depleted cells of endogenous HSP27 and ectopically expressed either a wild type or one of two mutant HSP27 alleles. HSP27-TriD has serine-to-asparagine phosphomimetic mutations at each of the three MK2 target serine residues. If HSP27 phosphorylation drives SASP mRNA stability, we reasoned that expression of HSP27-TriD in pre-senescent cells would create a senescent-like mRNA stability phenotype. Conversely, expression of HSP27-TriA, which has serine-to-alanine mutations at the key serine residues that prevent it from being phosphorylated, would result in a pre-senescent-like, lower mRNA stability state in senescent cells.

To assess the impact of HSP27 phosphorylation on mRNA stability in senescent cells, we first simultaneously knocked down the endogenous HSP27 with an HSP27-directed short hairpin (shHSP27) and ectopically expressed shRNA-resistant FLAG-tagged HSP27 alleles in BJ fibroblasts. As previously reported in monocytes, HSP27-TriD expression is very low compared to HSP27-WT or HSP27-TriA [23], however in contrast to that report, AUF1 protein levels were not affected by the expression of these mutants in our system (Fig 4A). Untreated cells expressing HSP27-TriD had increased levels of SA-β-Gal staining, suggesting HSP27-TriD expression alone may promote senescence induction (Fig 4B). However, the expression of neither wild type nor HSP27 mutants affected the ability of cells to enter senescence following bleomycin treatment as quantified by significant increases in SA-β-Gal staining.

**Fig 4.**
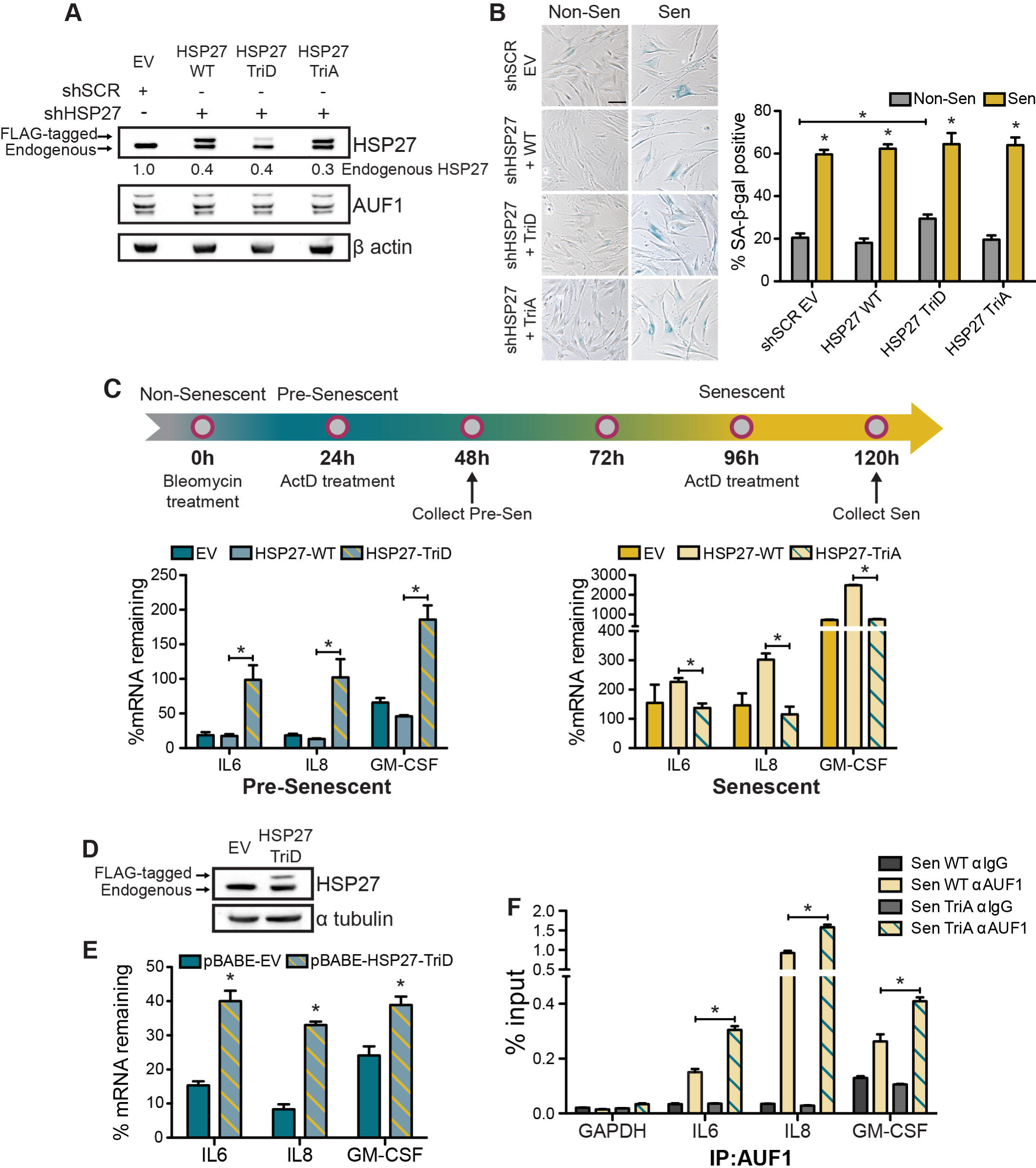
p38MAPK-dependent phosphorylation of HSP27 regulates SASP factor mRNA stability in senescent cells. (A) Immunoblot of simultaneous knockdown of endogenous HSP27 (lower band) and ectopic expression of FLAG-tagged HSP27 mutant alleles (upper band) in BJ fibroblasts. Percent of HSP27 knockdown after normalization to β-actin is presented under the HSP27 immunoblot. EV, empty vector. (B) Representative images and quantification of SA-β-Gal expression in non-senescent and senescent cells expressing pRESQ-shSCR-EV or pRESQ-shHSP27-Flag-HSP27 WT, TriD, or TriA. Images acquired with a 10x objective, scale bar = 100μm, n=2. (C) (Top) Timeline of senescence induction and ActD treatment. BJ fibroblasts expressing the HSP27 constructs were treated with bleomycin for 24h and ActD for 24h either immediately after bleomycin treatment (pre-senescent, left) or 96h post-bleomycin treatment (senescent, right), collected, and mRNA levels were analyzed by qRT-PCR. %mRNA remaining was calculated as the level of mRNA in ActD-treated cells relative to vehicle-treated cells, normalized to GAPDH. Representative experiment, n=3. (D) Immunoblot of HSP27 expression in cells transduced with pBABE-EV or pBABE-HSP27-TriD. (E) Cells expressing either pBABE-EV or HSP27 TriD were treated as in A and collected at the pre-senescent timepoint. Representative experiment, n=2, *=p<0.05. (F) BJ fibroblasts expressing a hairpin targeting HSP27 while simultaneously expressing flag-tagged HSP27 WT or TriA were induced to senesce by bleomycin treatment. Cells were collected 96h post-bleomycin (senescent) and anti-AUF1 RIP was performed using nonspecific IgG or an anti-AUF1 specific antibody. Representative experiment, n=3. *=p<0.05, error bars are ± S.E.M.

Cells expressing a control hairpin (shSCR) or shHSP27 with ectopic expression of HSP27-WT, HSP27-TriD, or HSP27-TriA were treated with bleomycin to induce senescence for 24h and then treated with ActD to inhibit transcription either immediately following bleomycin treatment (pre-senescent) or 96h post-bleomycin treatment (senescent). As expected, transcriptional inhibition in pre-senescent cells expressing HSP27-WT resulted in degradation of SASP factor mRNAs (Fig 4C, left). However, expression of the HSP27-TriD mutant resulted in significantly increased mRNA stability compared to cells expressing HSP27-WT, demonstrating that phosphorylated HSP27 protects SASP factor mRNAs from degradation and is sufficient to create a senescent-like mRNA stabilization state in pre-senescent cells (Fig 4C, left). Conversely, expression of the non-phosphorylatable mutant, HSP27-TriA, in senescent cells resulted in significantly decreased mRNA stability compared to HSP27-WT (4C, right). This demonstrated that HSP27 phosphorylation on p38MAPK-MK2 target sites is necessary for senescence-associated mRNA stabilization, and preventing this phosphorylation can destabilize mRNAs in senescent cells. Ectopic HSP27-TriD expression, albeit at low levels, recapitulates the mRNA stability phenotype we observed in shHSP27 expressing cells. To ensure that it was due to HSP27-TriD expression and not simply loss of endogenous HSP27 protein levels, we transduced FLAG-tagged HSP27-TriD without the hairpin into BJ fibroblasts and repeated the mRNA stability assay in pre-senescent cells (Fig 4D). We again found that ectopic expression of HSP27-TriD alone significantly increased the stability of IL6 and IL8 mRNAs, recapitulating a senescent-like mRNA stability state in pre-senescent cells (Fig 4E).

We previously found that AUF1 binding to SASP factor mRNAs decreases from pre-senescence to senescence in a p38MAPK-dependent manner [10]. Thus, we next wanted to determine whether the pre-senescent-like state created by expression of HSP27-TriA in senescent cells recapitulated the pre-senescent AUF1 binding state, *i.e.* increased AUF1 occupancy. We performed a RIP in senescent BJ fibroblasts expressing shHSP27 and either HSP27-WT or HSP27-TriA. When AUF1 was immunoprecipitated from these cells, we found that expression of HSP27-TriA significantly increased AUF1 binding from 0.15% to 0.30% of input for IL6, 0.92% to 1.58% of input for IL8, and 0.26% to 0.41% of input for GM-CSF mRNAs as compared to HSP27-WT (Fig 4F). Previous studies and our data from shHSP27-expressing cells (Fig 1F, G) argue that AUF1 binding to target mRNAs is not the only determinant of ARE-mediated mRNA degradation. However, we did find that AUF1 binding in HSP27-TriA-expressing senescent cells was increased while mRNA stability was decreased, recapitulating what is observed in wild type pre-senescent cells thus confirming that decreased AUF1 binding is a component of SASP mRNA stability regulation in senescent cells.

### The p38MAPK-MK2-HSP27 pathway promotes stromal-supported tumor cell growth

Previously we demonstrated that senescent stromal cells promote tumor cell growth, and inhibiting p38MAPK signaling in stromal cells significantly reduced this growth promotion [10]. Because p38MAPK inhibition studies in patients have suggested that cells develop resistance mechanisms to p38MAPK inhibition [33–35], and we find that MK2 is an important player in SASP mRNA stability, we investigated whether targeting the MK2 arm of the p38MAPK pathway would inhibit stromal-supported tumor growth. Along with the p38MAPK inhibitor CDD-111 (p38i′), we utilized CDD-450 (MK2i), which prevents p38MAPK from phosphorylating MK2, providing a more selective inhibition of the MK2 pathway while preserving alternate p38MAPK activity (Fig 2). Both p38i′ and MK2i inhibited the pathway as evidenced by reduced HSP27 phosphorylation. Furthermore, IL6 and IL8 upregulation and mRNA stabilization were significantly inhibited by treatment with both p38i′ and MK2i (Fig 5A-C). To test whether MK2 pathway inhibition prevented senescent stroma from promoting tumor growth, we co-cultured BJ skin fibroblasts with preneoplastic HaCaT keratinocytes expressing click beetle red (CBR) luciferase (HaCaT-CBR) as previously described, utilizing CBR activity as a direct measure of preneoplastic cell growth [6]. BJ fibroblasts were treated with bleomycin and p38i′ or MK2i for 96h to allow the fibroblasts to senesce, and then HaCaT-CBR cells were plated as indicated (Fig 5D). As expected, senescent BJ fibroblasts significantly increased the growth of HaCaT-CBR cells compared to non-senescent BJ fibroblasts (Fig 5E). In confirmation of our previous findings, p38MAPK inhibition abrogated this growth, by an average of 66.1%. Importantly, treatment with MK2i reduced tumor cell growth by 73.6% (Fig 5E) and this reduction in HaCaT-CBR growth was due to inhibition of the p38MAPK-MK2 pathway in the stroma specifically, because neither p38i′ nor MK2i treatment significantly affected the growth of HaCaT-CBR cells cultured independently.

**Fig 5.**
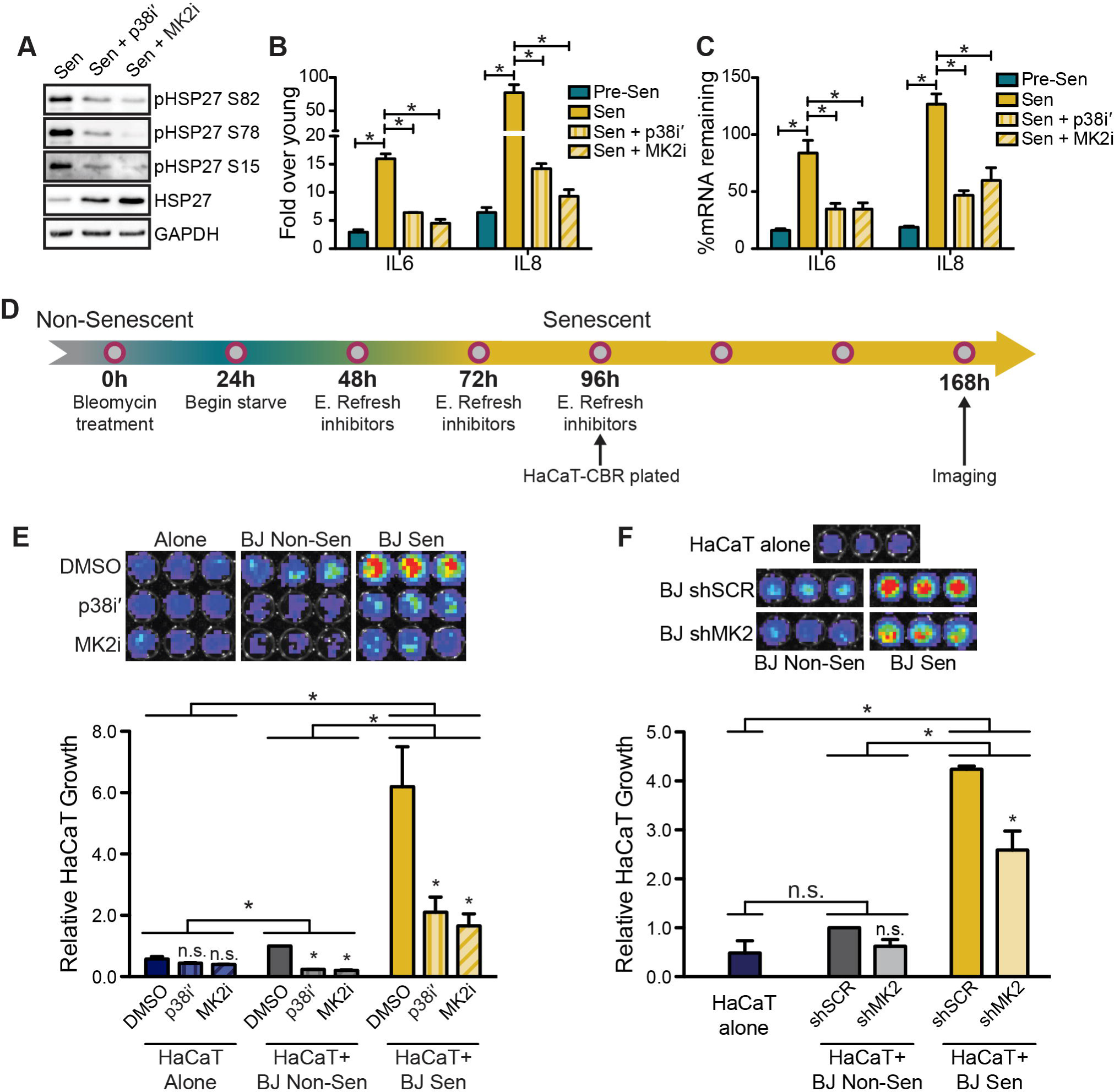
p38-MK2-HSP27 pathway activation in senescent stroma promotes tumor cell growth. (A) Immunoblot of HSP27 phosphorylation in cells treated as in Fig 3A, with p38i (CDD-111, 1μM) or MK2i (CDD-450, 1μM), respectively. (B) Upregulation and (C) mRNA stability of SASP factors in cells treated as in Fig 3A. SASP mRNA levels were quantified by qRT-PCR. Representative experiments, n=4 for B and n=2 for C. (D) Schematic of BJ fibroblast senescence induction, inhibitor treatment (for 5E), and plating of HaCaT-CBR cells for E and F. (E) BJ fibroblasts were plated in a 96-well plate and induced to senesce by 24h of bleomycin treatment. 24h after removal of bleomycin, cells were treated with DMSO, p38i′ or MK2i for 48h, refreshing the inhibitors after 24h. Once the fibroblasts were senescent 96h post-bleomycin treatment, click beetle red luciferase (CBR)-expressing HaCaT keratinocytes were plated either alone or on top of the non-senescent or senescent fibroblasts, and treated with DMSO, p38i′ or MK2i for E. To prevent disruption of the epithelial layer, inhibitors were not refreshed after HaCaT-CBR addition. HaCaT-CBR cell growth was measured 3 days after plating by addition of D-luciferin and bioluminesce quantification. Quantification normalized to HaCaT-CBR plated with Non-Sen, DMSO-treated BJ fibroblasts. Color bar: minimum=2434.5 flux-photons, maximum=10^5^ flux-photons. n=3. (F) Cells were treated as in E, without inhibitor treatment. HaCaT-CBR cells were plated onto either non-senescent on senescent BJ fibroblasts expressing either shSCR or shMK2. Quantification normalized to HaCaT-CBR plated with Non-Sen shSCR fibroblasts. Color bar: minimum=7000 flux-photons, maximum=2.2e10^5^ flux photons, n=2. n.s. = not significant, *=p<0.05, error bars are ± S.E.M.

To verify that the inhibition of HaCaT-CBR growth was due to stromal-specific p38MAPK-MK2 inhibition, we cultured HaCaT-CBR cells with senescent fibroblasts expressing either a control shSCR hairpin or on specifically targeting MK2 (shMK2). By utilizing MK2-depleted fibroblasts, we could ensure that any differences in the ability of the senescent fibroblasts to promote HaCaT-CBR cell growth was due changes in the stromal component of the co-culture, specifically MK2 activity, and not to effects of the inhibitors on the HaCaT-CBR cells. As before, the presence of senescent fibroblasts significantly promoted HaCaT-CBR cell growth relative to HaCaT-CBR cells grown alone or with non-senescent fibroblasts (Fig 5F). Although both shSCR- and shMK2-expressing senescent BJ fibroblasts significantly promoted HaCaT-CBR cell growth, there was a significant reduction in this growth promotion upon MK2 depletion. Therefore, the inability of MK2i-treated senescent fibroblasts to promote preneoplastic cell growth is due specifically to inhibition of MK2 in the stromal compartment rather than in the preneoplastic cells. Thus, MK2 activity in the stroma is required for the observed growth promotion. This reflects our previous findings that p38i’ treatment abrogates senescent stromal-driven preneoplastic cell growth and demonstrates that MK2 is the important downstream target of p38MAPK [10]. Taken together, these data indicate that MK2 pathway inhibition is a viable therapeutic strategy that can specifically prevent senescent stromal promotion of tumor cell growth.

## Discussion

The SASP is regulated by a multitude of signaling pathways at the levels of transcription, mRNA stability, and translation. In cells irradiated to induce senescence, p38MAPK is activated with slow kinetics, with peak phosphorylation of p38MAPK and HSP27 not occurring until 8-10 days post-irradiation [11]. Inhibition of p38MAPK with SB203580 or knockdown using an shp38α hairpin prevented the expression of SASP factors at the protein and mRNA levels, suggesting that p38MAPK regulates SASP factor transcription [10, 11]. Indeed, NFκB binding activity increases slowly in senescence reflecting p38MAPK activation, and p38MAPK activity is sufficient to induce NFκB binding.

In addition to transcriptional regulation, we previously demonstrated that p38MAPK regulates SASP factor expression through stabilization of target mRNAs in senescent cells, in an AUF1-dependent manner [10]. Here, we demonstrate that the presence of HSP27 and its p38MAPK-MK2-dependent phosphorylation state regulate SASP mRNA stability in pre-senescent and senescent cells. We find that increased levels of non-phosphorylated HSP27 in pre-senescent cells relative to senescent cells (Fig 3B), promotes SASP factor mRNA degradation, independently of AUF1 binding, as depletion of HSP27 increased both mRNA stability and AUF1 binding to SASP mRNAs (Fig 1G). Additionally, the stability of SASP factor mRNAs is dependent on the phosphorylation state of MK2 target sites on HSP27 (Fig 3G, 4C, and 5A-C). In pre-senescent cells, mRNA degradation correlates with reduced levels of HSP27 phosphorylation, while in senescent cells, increased mRNA stability correlates with increased HSP27 phosphorylation. The importance of HSP27 phosphorylation was underscored by our finding that expression of the phosphomimic HSP27-TriD in pre-senescent cells recapitulated the senescent state (i.e. stabilized mRNA). In contrast, expression of the phospho-dead HSP27-TriA resulted in destabilization of SASP mRNAs and thus created a pre-senescent-like state in senescent cells (Fig 4C). These data support the hypothesis that HSP27 phosphorylation is required to stabilize SASP mRNAs. Furthermore, phosphorylation of HSP27 is required for removal of AUF1 from SASP mRNAs in senescent cells, since HSP27-TriA expression increased AUF1 binding in senescent cells, leading to decreased SASP mRNA stability (Fig 4F). This suggests that in senescent cells, AUF1 binding and the phosphorylated state of HSP27 together regulate mRNA stability in senescent cells. A previous study in lymphocytes demonstrated that HSP27’s phosphorylation state regulates the abundance and stability of AUF1 resulting in mRNA stabilization in that system [21, 23]. However, we did not observe consistent changes in AUF1 levels upon HSP27 knockdown or expression of the HSP27 mutants, suggesting that an alternate p38-MK2-HSP27 dependent mechanism of regulating AUF1-mediated mRNA degradation is active in senescent fibroblasts.

AUF1 activity is regulated at several levels in different cellular contexts. In certain systems, AUF1-mediated mRNA degradation is regulated by AUF1 binding to target mRNAs, which can be affected by altering AUF1’s binding ability or through modulation of AUF1 levels within the cell. However, AUF1 binding alone is not always predictive of mRNA degradation, since AUF1 binding has also been reported to protect mRNAs from degradation [16, 18, 36]. Indeed, all four isoforms of AUF1 can promote mRNA degradation and/or protection depending on cell type and context. The isoforms have different ARE-binding affinities, suggesting that AUF1 isoforms control stability through differential binding of specific AREs [13, 15, 17, 18, 37]. Furthermore, AUF1 activity can be regulated by post-translational modifications. In THP-1 lymphocytes, phosphorylated p40^AUF1^ is associated with TNFα mRNA degradation whereas non-phosphorylated p40^AUF1^ is associated with longer TNFα mRNA half-life and higher levels of translation, suggesting phosphorylated p40^AUF1^ is actively involved in AMD [19, 20]. Conversely, phosphorylation of p40^AUF1^ is required for ubiquitin-mediated AUF1 degradation in HeLa cells, resulting in increased mRNA stability [21]. Together, these observations illustrate the complexity and context-specificity of the regulatory mechanisms governing AUF1’s roles in mRNA stability.

Our data suggest that the phosphorylation state of HSP27 and the binding of AUF1 impact mRNA stability in pre-senescent and senescent cells (Fig 6). Indeed, non-phosphorylated HSP27 favors AMD as evidenced by our findings that (i) in pre-senescent cells, which have low p38MAPK activity and reduced levels of phosphorylated HSP27, HSP27 depletion results in mRNA stabilization despite increased AUF1 binding, and (ii) expression of the non-phosphorylatable allele HSP27-TriA results in mRNA degradation in senescent cells. Thus, our data suggest an important role for non-phosphorylated HSP27 in AMD in pre-senescent cells, because if phosphorylated HSP27 were the only state in which HSP27 regulated mRNA stability, then HSP27 depletion in pre-senescent cells should result in increased mRNA degradation due to the lack of basal phospho-HSP27 in the system. This activity may occur through HSP27-dependent recruitment or activation of other ASTRC members or other ARE binding proteins.

**Fig 6.**
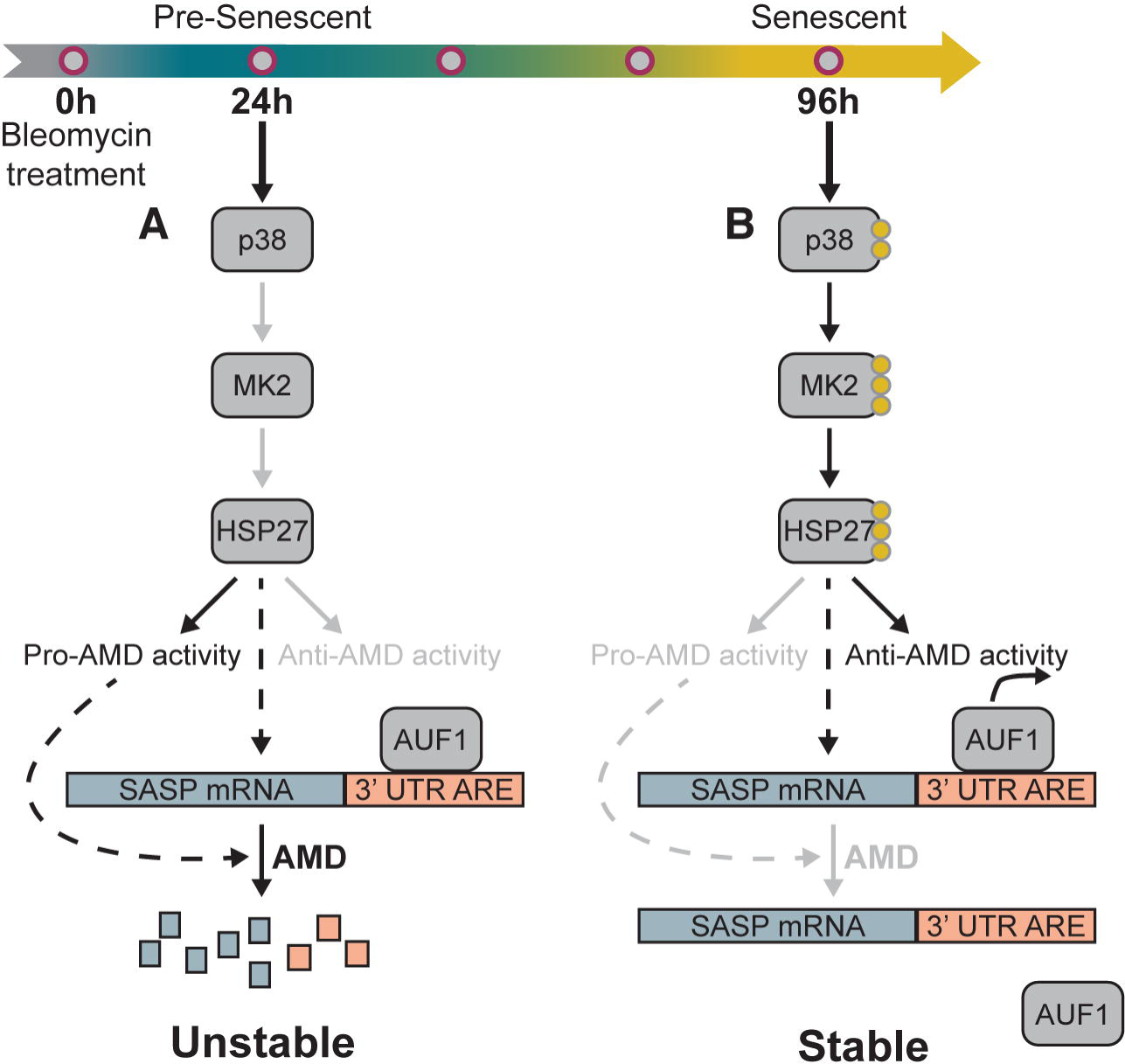
Model of p38MAPK-MK2-HSP27-AUF1 pathway regulation of SASP factor mRNA stability in pre-senescent and senescent fibroblasts. (A) In pre-senescent cells, p38MAPK is not yet activated to the level observed in senescent cells, so p38MAPK-MK2-dependent phosphorylation of HSP27 is low and AUF1 binds target mRNAs. Non-phosphorylated HSP27 performs a putative pro-AMD activity, and the levels of phospho-HSP27 are insufficient to abrogate this function and thus mRNA degradation is dominant. (B) In senescent cells, p38MAPK phosphorylation activates the MK2-HSP27 pathway stabilizing SASP mRNAs, potentially through several mechanisms. Reduced levels of non-phosphorylated HSP27 result in decreased HSP27-dependent pro-AMD activity, and phosphorylation of HSP27 on MK2 target sites results in AUF1 removal and stabilization of target SASP mRNAs.

Together, these observations suggest that senescence-associated AMD is regulated by HSP27 and AUF1 activity, and that the role of HSP27 may be upstream of AUF1 in the pre-senescent state. In pre-senescent cells, p38MAPK activity is low, and both AUF1 and HSP27 actively promote AMD, resulting in SASP mRNA degradation (Fig 1C, 6A). This suggests that non-phospho-HSP27 is its AMD-promoting state. When HSP27 is depleted in pre-senescent cells, the lack of non-phosphorylated, AMD-promoting HSP27 results in mRNA stabilization despite AUF1 binding to target mRNAs (the increase in AUF1 binding observed in shHSP27 cells may be due to the reduction of basal levels of phosphorylated, or AMD-inhibiting, HSP27). This indicates that although AUF1 is necessary, AUF1 binding is insufficient to drive AMD without HSP27 in pre-senescent cells. Once the p38MAPK pathway is activated and cells are senescent (Fig 6B), AUF1 activity appears to be regulated at the level of binding, since both inhibition of the p38MAPK pathway as well as expression of the non-phosphorylatable HSP27-TriA result in increased AUF1 binding and subsequent destabilization of SASP mRNAs (Fig 4F, [10]).

Like p38MAPK inhibition, MK2 depletion and MK2i treatment prevent HSP27 phosphorylation and SASP mRNA stabilization, and our data from HSP27-TriA and HSP27-TriD suggest that phosphorylation of MK2 target sites on HSP27 drives this phenotype. Recently, Herranz *et al.* demonstrated that mTOR modulates MK2 translation in the context of oncogene-induced senescence (OIS), allowing MK2 to phosphorylate and inactivate ZFP36L1, another mRNA destabilizing protein that binds AU rich elements similarly to AUF1. They suggest that this ZFP26L1 phosphorylation then prevents ARE-containing SASP mRNAs from being degraded in senescent cells resulting in SASP factor upregulation. Our previous data demonstrate that AUF1 binds and targets SASP mRNAs for degradation in pre-senescent but not senescent cells and that depletion of AUF1 from pre-senescent cells stabilizes SASP mRNAs [10]. Here we demonstrate HSP27 regulates AUF1 occupancy on SASP mRNAs in senescent cells in an MK2-dependent manner. These findings together suggest that MK2 has a multifaceted role in the regulation of SASP factor mRNA stability and upregulation, further supporting MK2 as a promising drug target for inhibition of the p38MAPK pathway.

The p38MAPK pathway regulates the expression of many pro-inflammatory cytokines and other factors responsible for chronic inflammatory diseases such as chronic obstructive pulmonary disease (COPD), rheumatoid arthritis (RA), psoriasis, and Chron’s disease [35], and therefore has been an attractive target for treating the inflammation associated with these diseases. However, clinical trials for some of these diseases have so far been disappointing, due to problems such as low efficacy, adverse side effects, and a rebound in levels of inflammatory cytokines [33, 38, 39]. This rebound effect may be due to inhibition of downstream anti-inflammatory substrates of p38MAPK (e.g. MSK1/2 or MKP1). Because we have found that p38MAPK inhibition within a senescent microenvironment prevents tumor growth [10], these findings raise questions about the potential efficacy of p38MAPK directed therapies. MK2 may provide an alternative, more specific therapeutic target that could prove to be more effective.

Current cancer treatment modalities largely target tumor cells, neglecting the critical impact of the tumor microenvironment on tumor establishment and growth. Modulation of the tumor microenvironment is sufficient to promote or restrain tumor cell growth, demonstrating that stromal-specific therapies or co-therapies are a promising therapeutic avenue, and targeting the SASP through p38MAPK and MK2 activity is a possible means to do so. Herranz *et al.* demonstrated that the mTOR/MK2/ZFP36L1 pathway mediates the pro-tumorigenic aspects of the SASP. Here, we have demonstrated that MK2 also regulates the SASP through HSP27 and AUF1. Furthermore, we have shown that inhibition of the MK2 pathway in the stromal compartment is as effective as p38MAPK inhibition at limiting the expression and mRNA stabilization of SASP factors, thereby preventing senescent stroma from promoting the growth of pre-neoplastic HaCaT keratinocytes *in vitro* and demonstrating that targeting the MK2 pathway is a viable means of circumventing the putative shortcomings seen with p38MAPK inhibition in a clinical setting. Further studies in an *in vivo* setting will investigate the viability of inhibiting the MK2 pathway by CDD-450 as a stromal-specific cancer therapy.

## Supporting information

Supp Table Antibodies

## Acknowledgements

We thank Julie Prior and the ICCE Institute at Washington University School of Medicine for live cell imaging, and Megan Ruhland and Kevin Flanagan for helpful comments. This work was supported by the Cancer Biology Pathway, Siteman Cancer Center at Barnes Jewish Hospital and Washington University Medical School (HM), NIH grants GM007067 to (EA) and CA130919 (SAS), and American Cancer Society Research Scholar Award (SAS).

## Supporting Information

**S1 Table.** Extended antibody information for primary antibodies used.

**S1 Fig. Full western blot membranes used for Fig 1.** Images acquired using Bio-Rad Chemidoc XRS+ imager and ImageLab software. Images were unaltered in ImageLab and copied directly. Due to the differences in acquisition times, ladder markings are not always visible. (A) Complete membranes for Fig 1B. * = top of membrane cut off during acquisition. Multiple bands of unknown origin were observed for all antibodies utilized, but the darkest band was always present at the expected molecular weight. (B) Complete membranes/images used for Fig 1D. AUF1 and total HSP27 were imaged together.

**S2 Fig. Full western blot membranes used for Fig 3.** Images acquired using Bio-Rad Chemidoc XRS+ imager and ImageLab software. (A) Membranes used for Fig 3B. Lower lane in β actin blot is nonspecific band sometimes observed. (B) Membranes used for Fig 3E. Top row, left = merge of brightfield image of ladder and chemiluminescence image shown on left. Nonspecific lower bands in phosphorylated HSP27 blots are occasionally observed when higher concentrations of whole cell lysate are loaded, and may originate form degradation products of HSP27.

**S3 Fig. Full western blot membranes used for Fig 4.** Images acquired using Bio-Rad Chemidoc XRS+ imager and ImageLab software. (A) Membrane images used for Fig 4A. AUF1 and total HSP27 were imaged together. (B) Membrane images used for Fig 4D. Total HSP27 and α tubulin were imaged together.

**S4 Fig. Full western blot membranes used for Fig 5.** Images acquired using Bio-Rad Chemidoc XRS+ imager and ImageLab software. Membrane images used for Fig 5A.

