## Supplementary material for "The p38MAPK-MK2-HSP27 Pathway Regulates the mRNA Stability of the Senescence-Associated Secretory Phenotype": Supp Table Antibodies

| **Target** | **Dilution** | **Species** | **Clonality** | **Clone** | **Antibody registry** | **Catalog number** | **Lot number** | **Company** | **Immunegen** | **Validation** |
| --- | --- | --- | --- | --- | --- | --- | --- | --- | --- | --- |
| **pHSP27 Ser15** | 1:1000 | Rabbit | Monoclonal |  | AB_330738 | 2404S | 4 | Abcam, Cambridge, MA | Synthetic phospho-peptide of residues surrounding Ser15 of human HSP27 | Decrease in band intensity from cells expressing shHSP27 hairpin |
| **pHSP27 Ser78** | 1:2000 | Rabbit | Monoclonal | Y175 | AB_733030 | ab32501 | GR102949-2 | Cell Signaling Technologies, Danvers, MA | Synthetic phospho-peptide of residues surrounding Ser78 of human HSP27 | Decrease in band intensity from cells expressing shHSP27 hairpin |
| **pHSP27 Ser82** | 1:1000 | Rabbit | Polyclonal |  | AB_331644 | 2401S | 1 | Cell Signaling Technologies, Danvers, MA | Synthetic phospho-peptide of residues surrounding Ser82 of human HSP27 | Decrease in band intensity from cells expressing shHSP27 hairpin |
| **total HSP27** | 1:1500 | Mouse | Monoclonal | G31 | AB_10696744 | 2402S | 8 | Cell Signaling Technologies, Danvers, MA | Full length HSP27 (species not specified) | Decrease in band intensity from cells expressing shHSP27 hairpin |
| **MAPKAPK2** | 1:1000 | Rabbit | Polyclonal |  | AB_10827910 | 3042P | 3 | Cell Signaling Technologies, Danvers, MA | C-terminal region of human MAPKAPK2. | Decrease in band intensity from cells expressing shMK2 hairpin |
| **pp38 Thr180/Tyr182** | 1:1000 | Rabbit | Polyclonal |  | AB_2492195 | p190-1802 | cs1011r | PhosphoSolutions, Aurora, CO | From Rat p38MAPK, residues surrounding the phospho-Thr180 and Tyr182 | PhosphoSolutions states immunolabeling is blocked by preadsorption with the antigen phosphopeptide, but not the non-phosphopeptide of the same region |
| **total p38α** | 1:1000 | Rabbit | Polyclonal |  | AB_10694846 | 9218S | 5 | Cell Signaling Technologies, Danvers, MA | C-terminal region of human p38α | Decrease in band intensity from cells expressing shp38α hairpin [10] |
| **AUF1** | 1:4000 | Rabbit | Polyclonal |  | AB_2117338 | 07-260 | 2646922 | EMD Millipore, Billerica, MA | Purified human AUF1 | Millipore states antibody has been validated by WB. Decrease in band intensity from cells expressing shAUF1 hairpin [10] |
| **α-tubulin** | 1:5000 | Rat | Monoclonal | YL1/2 | AB_305328 | ab6160 | GR57687-1 | Abcam, Cambridge, MA | Full length *S. cerevisiae* native protein |  |
| **γ-actin** | 1:5000 | Rabbit | Polyclonal |  | AB_10003448 | NB600-533 | A2 | Novus Biologicals, Littleton, CO | Human Actin Gamma 1 N-terminus |  |
| **β-actin** | 1:2000 | Mouse | Monoclonal | AC-15 | AB_476692 | A1978 | 117K4873 | Sigma Aldrich, St. Louis, MO | Slightly modified, conjugated peptide corresponding to N-terminus of cytoplasmic β actin |  |
| **GAPDH** | 1:2500 | Rabbit | Monoclonal | GAPDH-71.1 | AB_1078991 | G8795 | 028K4859 | Sigma Aldrich, St. Louis, MO | Rabbit GAPDH |  |
