## Supplementary figures and images for "The p38MAPK-MK2-HSP27 Pathway Regulates the mRNA Stability of the Senescence-Associated Secretory Phenotype"

### Suppl fig 1

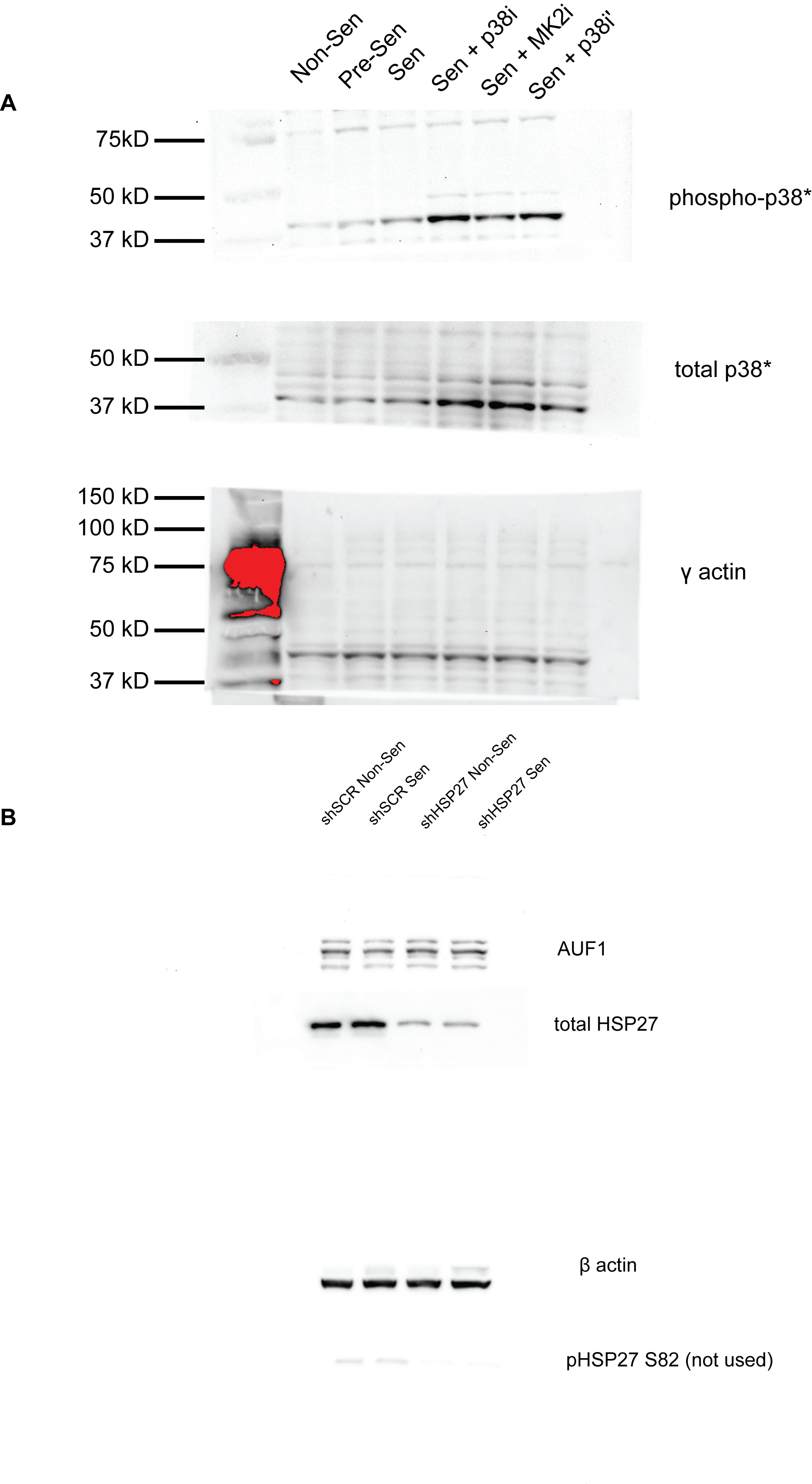

### Suppl fig 2

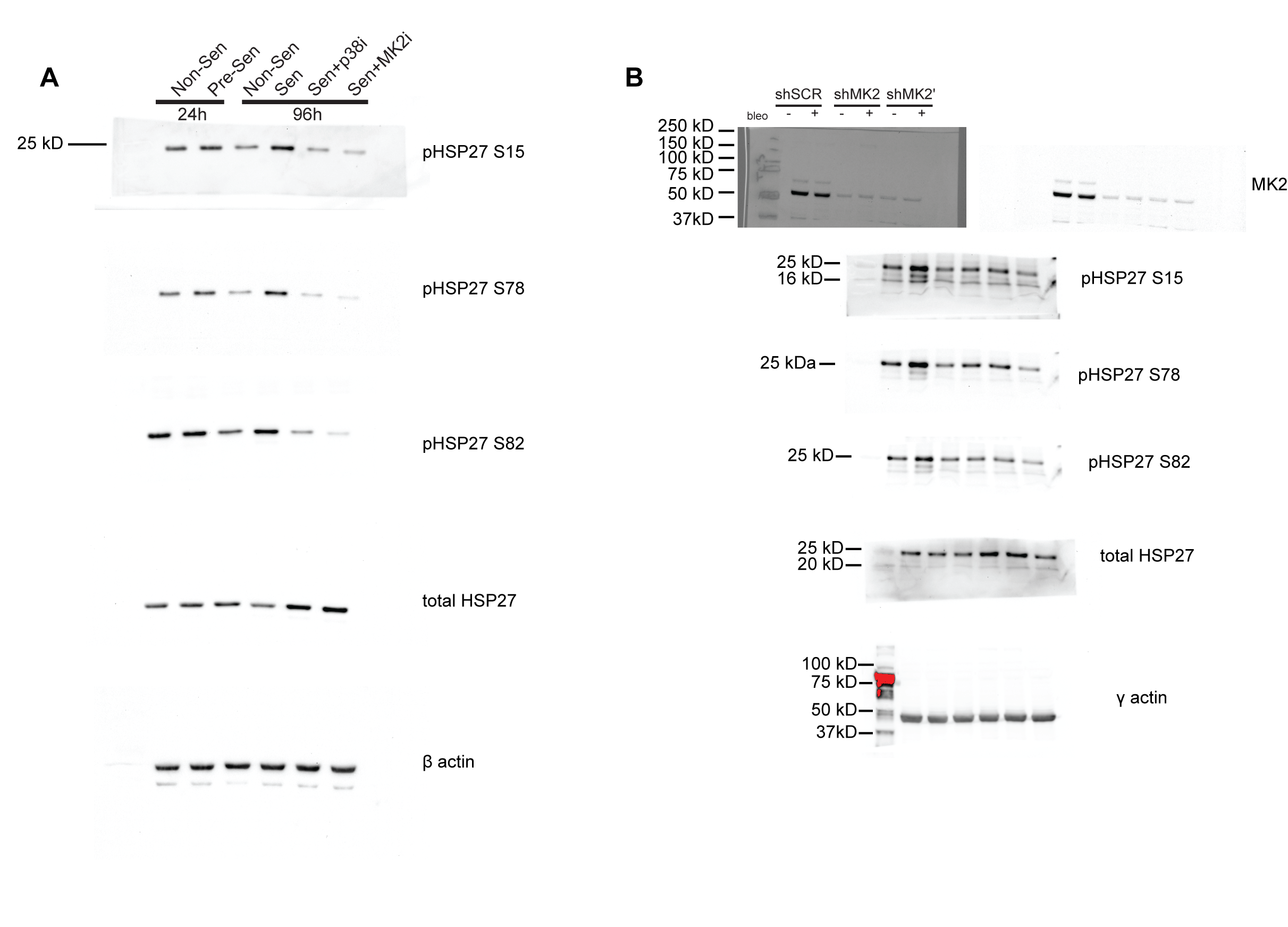

### Suppl fig 3

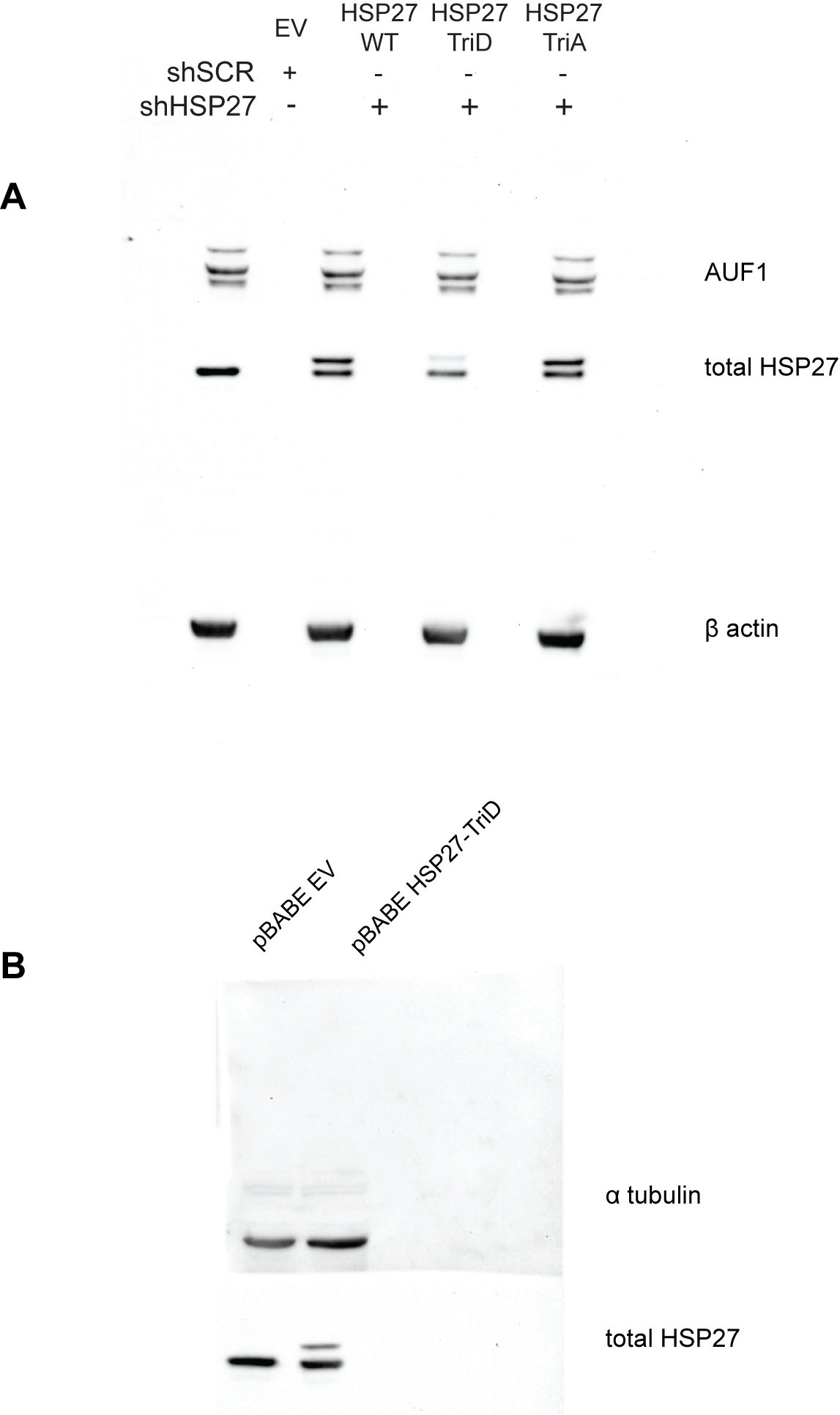

### Suppl fig 4

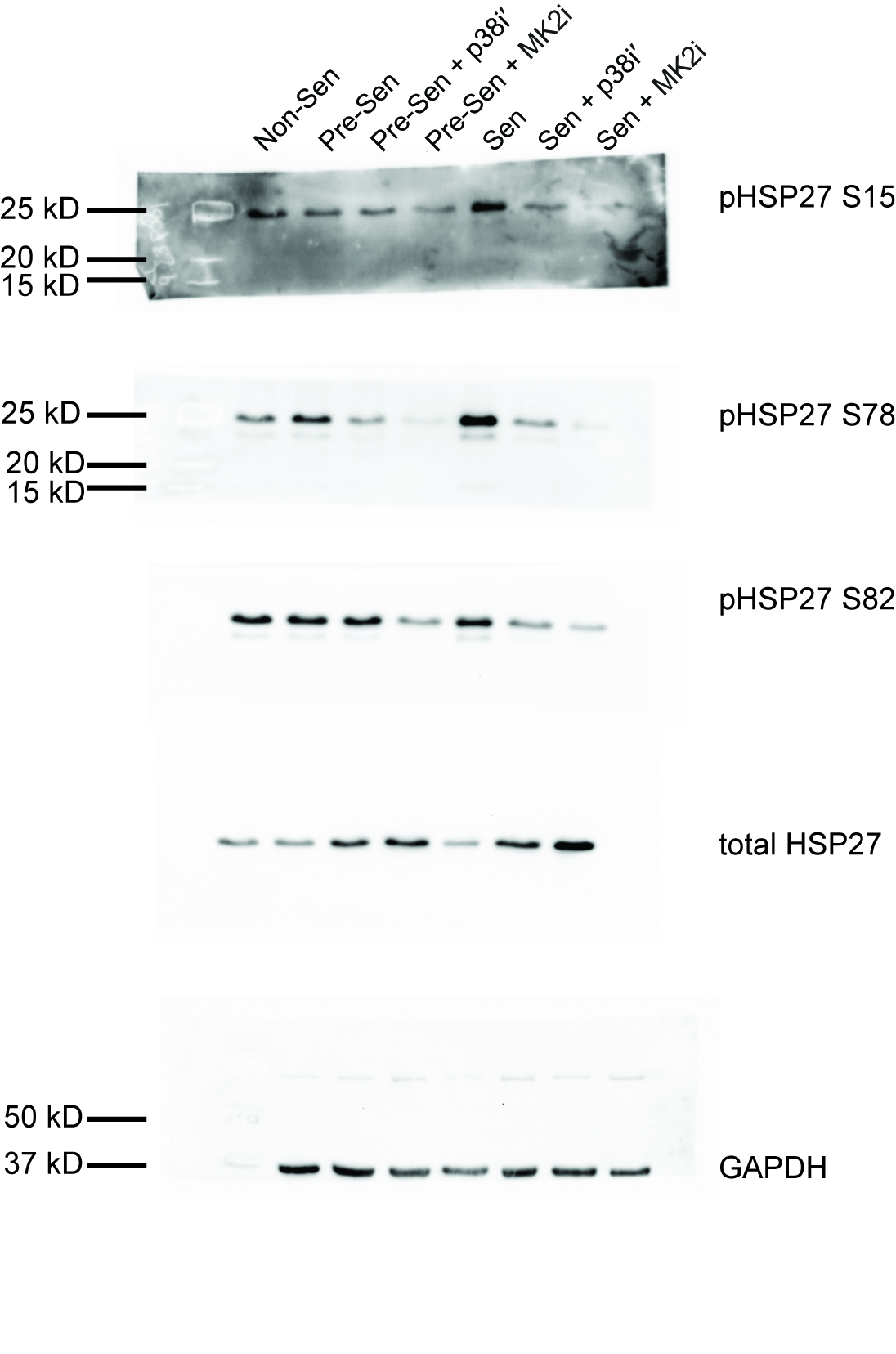
